# A mutation-mediated host shift drives Avian paramyxovirus type 1 evolution

**DOI:** 10.1101/2023.06.05.543675

**Authors:** Han Chang, Shengyong Feng, Xiaolu Tang, Ziming Wang, Wen Su, Shuyi Han, Guanghao Li, Bin Hu, Shengfan Jing, Bo Wang, Chen Xiang, Yanan Xing, Ye Wang, Jian Lu, Hongxuan He

## Abstract

PPMV-1, an antigenic variant of APMV-1, associated with specific pigeon host species. However, its evolutionary strategy and underlying drivers of host specificity remain unknown. In this study, we collect the outbreak data on a global scale to investigate its evolutionary dynamics, and provide an evidence-supported analysis the host shift of PPMV-1 from chickens to pigeons, and this shift is driven by the P protein. Our data indicated that the viruses in the United States and China have undergone convergent evolution. We find that three mutations of P protein, especially R163G, can significantly affect the adaptation of APMV-1 in pigeons. Mechanistically, sensor LSm14A inhibits the replication APMV-1 in DF-1 cells, and R163G substitutionon P protein increase LSm14A degradation. We propose the host shift drive the evolution of PPMV-1 and the underlying mechanism, offering new insights into the adaptive evolutionary process of the virus.

## Introduction

Many emerging viruses are the result of pathogens jumping from their original hosts to novel species. Successful virus emergence may occur through two distinct processes: host range expansion or shift. Range expansion allows the pathogen to infect more host species without altering its original gene pool (Thines 2019). Host shift, on the other hand, increases genetic differentiation in the pathogen gene pool, resulting in specialization to the novel host (Longdon et al, 2014; Thines 2019). Virus host shift involves different ecological and evolutionary processes. If the main factor leading to the emergence is ecological factors, and the shift does not require adaptation, the reasons for the host shift are called ecological drivers. Yet, if a virus emerges in a new host requiring genetic changes, the cause is known as an adaptive driver, although ecological drivers may also be present in this case (Pepin et al, 2010). The Avian paramyxovirus serotype 1 (APMV-1) of genus Avulavirus within the Paramyxoviridae family (Mayo 2002), has a negative-sense, single-stranded RNA genome approximately 15.2 kb in length that contains six genes in the order of 3’-NP-P-M-F-HN-L-5’. Two additional proteins, V and W, are generated by an RNA-editing event that occurs during the transcription of the P gene (Steward et al, 1993). The virus can infect a wide variety of Avian species (Wan et al, 2004). Pigeon paramyxovirus type 1 (PPMV-1), an antigenic variant of APMV-1 (Avian paramyxovirus type 1) (Collins et al, 1994; Mayo 2002; Ujvari et al, 2003), displays remarkable species-specificity to pigeons, is likely to be the result of APMV-1 host shift. RNA viruses appear to be particularly prone to host shifts, a mechanism that is a major driver of virus evolution in nature (Alkhamis et al, 2020; He et al, 2020). However, some studies have reported that PPMV-1 infections occurred in transplant patients under immunosuppressive therapy (Goebel et al, 2007; Kuiken et al, 2018). A recent study reports a case of severe pneumonia and eventual death in an immunocompetent patient (Zou et al, 2022). Has PPMV-1 evolved by host range expansion or shift, and the major evolutionary drivers of PPMV-1 and the underlying mechanism of PPMV-1 evolution have not been explained.

Adaptation generally is considered to affect the likelihood that a virus will be able to successfully emerge in a new host species and have high fitness (Pepin et al, 2010). Viral fitness is generally measured by replication capacity (Orr 2009). The polymerase complex, the minimal replication unit, of numerous viruses is related to viral tropism and host range (Mehle and Doudna 2009; Bortz et al, 2011; Bradel-Tretheway et al, 2011; Mehle et al, 2012; Long et al, 2019). The PPMV-1 polymerase complex composed of the P and L proteins, assembles with viral RNA and nucleoprotein (NP) to mediate transcription and replication of the viral genome. Serial passages of PPMV-1 in chickens result in host adaptation driven by mutations in the polymerase complex, suggesting that changes in this complex can drive host shifts (Dortmans et al, 2011; Olszewska-Tomczyk et al, 2018). Paramyxovirus P protein is essential for viral RNA synthesis and other biological processes (Lamb and Kolakofsky 1996). The V protein affects the host range of the virus via its species-specific IFN antagonist activity (Park et al, 2003). Therefore, the role of P protein in host shift appears to be particularly important, yet it has been rarely addressed in previous studies.

In the present study, we collect global data and used a phylogeographic Bayesian statistical framework to reconstruct PPMV-1 spatial spread over time and the virus transmission history among host species, and identify and characterize the cumulative molecular changes present in naturally occurring PPMV-1 that were responsible for the host adaptation. To further decipher its adaptation process, we evaluate the selection pressure of the six PPMV-1 proteins and genetic changes on P protein, explored the functional locus of P protein in this evolutionary process, and its underlying molecular mechanisms.

## Results

### Subtype and Host Changes during Evolution

PPMV-1 is an antigenic variant of APMV-1, and new subtypes are emerging (Liu et al, 2003; Chen et al, 2013). To confirm the main factor leading to the virus emergence is ecological or adaptive, we constructed a Bayesian tree based on complete F protein sequence using global isolates information to explore the epidemic genotypes of PPMV-1 in each country and the host composition. The tree was divided into two large clades at the root. One clade consisted of genotype XX and XXI, and the other was genotype VI. Genotype VI was further split into 8 subtypes. The topological structure of the phylogenetic tree was associated with the geographical distribution of these PPMV-1 strains. Some subtypes were predominant in particular countries, for example VI.2.2.2, VI.2.1.1.2.1, and VI.2.1.1.2.2 in China, and VI.2.1.2 and VI.2.1.1.1 in the United States (supplementary fig. 1A). We then analyzed transmission intensity using BSSVS to infer the forces that drive specific genotypes to dominance in different countries. The model identifies Iraq as the PPMV-1 distribution center, consistent with its previously identified origin in that region (Kaleta and Deursen 1985), and Europe to Africa spread is more frequent (supplementary fig. 1B). We further inferred the isolates from China or the United States formed special subtypes due to relatively infrequent country-to-country spread (supplementary fig. 1B). We also found that the host range of different genotypes was significantly different. Genotype XX mostly infects non-pigeon birds, and Genotype XXI and VI are primarily found in pigeons. The pigeon proportion as hosts of the United States and China subtype increases with time (fig. 1D and supplementary fig. 1C). The results indicated host shift may be the main evolutionary strategy of PPMV-1.

**Figure 1.**
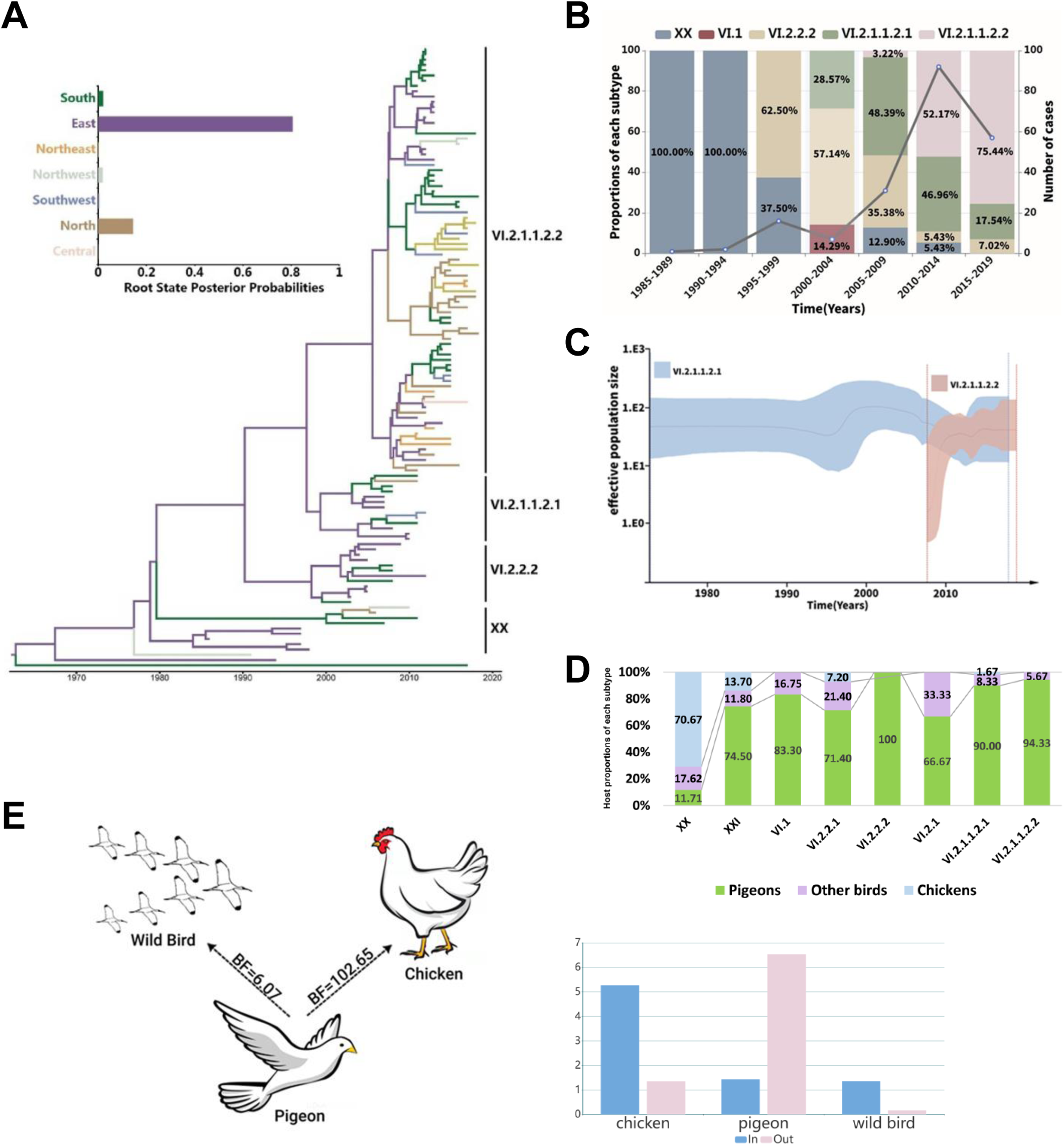
Host shift during evolution. **A.** Time-scaled maximum clade credibility F gene phylogeny with branches colored according regions. 34 provinces and cities were divided into seven regions on the basis of geographic proximity: Northwest (Xinjiang, XJ; Qinghai, QH; Gansu, GS; Ningxia, NX, Shaanxi, ShaX), North (Inner Mongolia, NM; Hebei, HeB; Shanxi, SX; Beijing, BJ; Tianjin, TJ), Northeast (Heilongjiang, HLJ; Jilin, JL, Liaoning, LN), East (Shandong, SD; Anhui, AH; Fujian, FJ; Zhejiang, ZJ; Jiangxi, JX; Jiangsu, JS; Taiwan, TW; Shanghai, SH), Southwest (Xizang, XZ; Yunnan, YN; Sichuan, SC, Guizhou, GZ; Chongqing, CQ), Central (Henan, HeN; Hubei, HuB; Hunan, HuN;), South (Guangdong, GD; Guangxi, GX; Hainan, HaiN; Aomen, AM; Hongkong, HK). **B.** The proportion of various subtypes between 1985 and 2019 in China. The number of cases is depicted by the line. **C.** Comparison of Bayesian skyline plot between subtype VI.2.1.1.2.1 and VI.2.1.1.2.2. The shaded red and blue bands give the 95% HPD intervals of the estimates for VI.2.1.1.2.1 and VI.2.1.1.2.2, respectively. **D.** Changes of host species distributions during the evolution in China. **E.** Significant transmission routes between host species inferred using the BSSVS approach. BF values of each route were on top of the arrows. The chart summarized total relative in and out transitions for each host.

Due to the special subtypes, we focus on the virus isolated in China. 241 PPMV-1 outbreaks in 26 of the 34 provinces, municipalities, and minority autonomous regions in China were recorded (Supplementary Table 4). PPMV-1 prevalence was highest in the eastern and southern regions, especially in Guangdong and Guangxi Provinces (supplementary fig. 2A). To create a geographically discrete partitioning scheme, 34 provinces and cities were divided into seven regions on the basis of geographic proximity as previously described (Bi et al, 2020). The prevalent subtypes in each of the seven regions are largely the same and have no regional preference. The PPMV-1 phylogenetic tree in China is ladder-like (that is the subtypes are replaced one by one) and the Bayesian analysis placed the root of the tree in the East, with a posterior probability of 0.81, which suggested that PPMV-1 emerged from East China (fig. 1A). In China, the regional distribution of PPMV-1 subtypes is relatively uniform, which is supported by BSSVS model analysis (supplementary fig. 2B). We also observed migration links from the South and the Northwest to the Northeast that are much stronger than from other regions (supplementary fig. 2B). The highest Bayes factors (BF, BF> 1000) was observed from the South to the East or the Southwest, from the East to the Center and the Northeast, and from the Northeast to the Center (supplementary fig. 2B). The highest migration rate (MR>= 1.5) is from the South and the Northwest to the Northeast. The general transmission trend is from the South to the North (supplementary fig. 2B).

The MCC tree of all isolates recorded in China revealed that XX was the original genotype, and was complete replaced from 2000 by a new variant (VI). Genotype Ⅵ emerged in 1992 in Zhejiang, and mainly the VI.2.2.2 subtype initially. Subsequently VI.2.1.1.2.1 appeared and increased rapidly. VI.2.1.1.2.2 replaced VI.2.1.1.2.1 as a dominant PPMV-1 subtype in the past ten years (fig. 1B). We observed the virus continues to spread rapidly, a dramatic rise in number of PPMV-1 cases from 2000-2014 and a decline in recent years (fig. 1B). Phylodynamic modeling of the effective population sizes of dominant subtypes (VI.2.1.1.2.1 and VI.2.1.1.2.2) in recent ten years indicated that VI.2.1.1.2.2 expanded rapidly since its emergence, and exceeded VI.2.1.1.2.1 in 2010 (fig. 1C), which is consistent with the percent of subtypes (fig. 1B).

Host range changes of each subtype in China were similar to the global trend, with the proportion of pigeons as hosts increases with time (fig. 1D). We next used a Bayesian stochastic search variable selection (BSSVS) procedure to identify PPMV-1 host shifts among pigeons, chickens, and wild birds. BF was used to estimate statistical support for these shifts. We identified two highly supported (BF > 3) host shifts from pigeons to chickens and wild birds. To quantify the magnitude of these host shifts, we estimated the number of host switching events (Markov jumps) per unit of time. Pigeons were the most frequent source and chickens were the most frequent recipients during host shift events. The virus had a capacity to spill over into non-pigeons but seldom came back (fig. 1E). We therefore confirmed that host shift may be the driving force of PPMV-1 evolution.

### Three P protein Amino Acid Substitutions Drive the PPMV-1 Evolution

In order to identify proteins driving the virus evolution, the average non-synonymous substitution/synonymous substitution rate was calculated for the whole protein. The data showed that all the six proteins were under negative selection, and dN/dS ratio of P and F protein are higher than the other proteins (fig. 2B). Notably, The dN/dS of P protein is about three times that of NP, M and L protein. What’ more, some regions cumulative dN/dS of the P protein and F protein are larger than 1 (fig. 2A), these proteins may become evidence of adaptive evolution.

**Figure 2.**
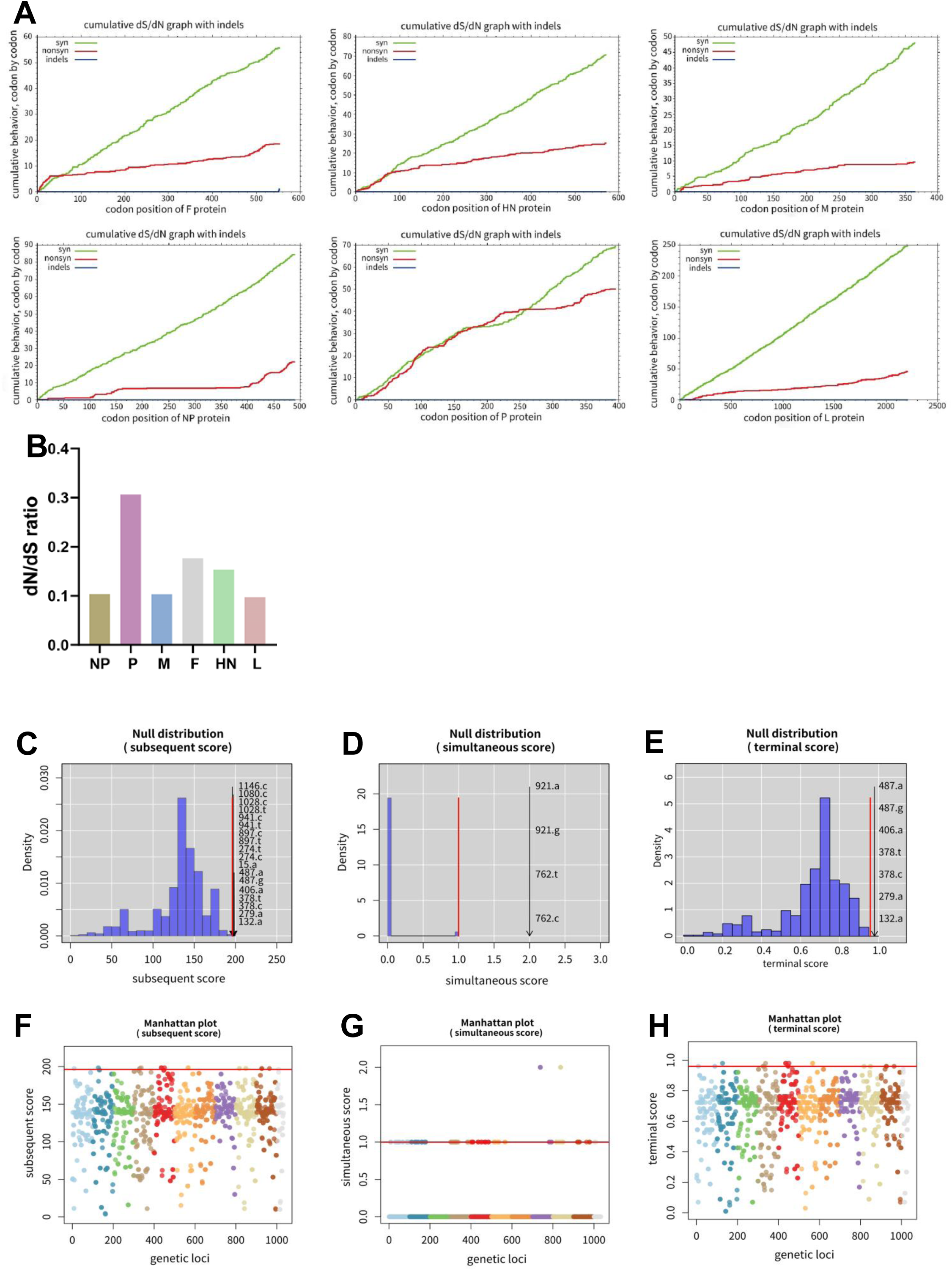
The potential mutations in P gene may be associated with host shift. **A.** Evolutionary rates are estimated by cumulative dS/dN. The regions under positive selection are indicated by black arrows. **B.** dN/dS ratio of the six whole proteins. **C-H.** Using treeWAS analysis, 18, 4 and 7 mutations were detected by **C.** subsequent score **D.** simultaneous score **E.** terminal score, respectively. The sites were labeled on the the corresponding figures. Manhattan plots for **F.** subsequent score **G.** simultaneous score **H.** terminal score showing association score values for P gene. The significance thresholds above the red line indicate significant associations.

To confirm the role of F and P protein in host shift, the evolution of PPMV-1 from different hosts was analyzed using Maximum-Likelihood trees and median-joining networks based on complete F (supplementary fig.3A and 3C) and P gene (Figures S3B and S3D) sequences. Obvious ‘host jumps’ (a cross-species transmission of a pathogen that can lead to successful infection) was observed in the phylogenetic analysis by both methods, and the amino acid of F and P protein evolutionary are positively correlated (supplementary fig. 3E)

Since virus adaptation is crucial to host jumping, if the genetic markers that adapt to new hosts can be identified and their impact on virus transmission can be determined, then the genome information can be used to predict future risks (Pepin et al, 2010). Some sites are host-specific and can be used as markers to distinguish hosts (Allison et al, 2014; Lee et al, 2019; Pascelli et al, 2020). We therefore explored which proteins and sites play an important role in host shift, and then used phylogenetic tree-based approach to genome-wide association studies (treeWAS) to make these inferences, and linking genotype to phenotype by testing for statistical associations between the two (Collins and Didelot 2018). The results showed that no nucleotide sites related to host species were found in F gene (supplementary fig. 3E), and 22 nucleotide sites (harbored 5 non-synonymous mutations: T93K, W136R, R163G, P314L, A343V) in P gene were found to be reliable genetic markers for host adaptation. (fig. 2C-H).

Since the emergence of APMV-1 was previously attributed to adaptation in chickens and other wild birds, it is now clear that the emergence of PPMV-1 involved the host shift event. The PPMV-1-specific residues were likely acquired during the virus evolution in pigeons. From the phylogenetic tree, the five mutations were clearly displayed during the evolution from APMV-1 into non-pigeon PPMV-1 till pigeon-specificity (fig. 3A). Interestingly, the amino acid substitutions occur largely in host shift rather than genotype switch (fig. 3A).

**Figure 3.**
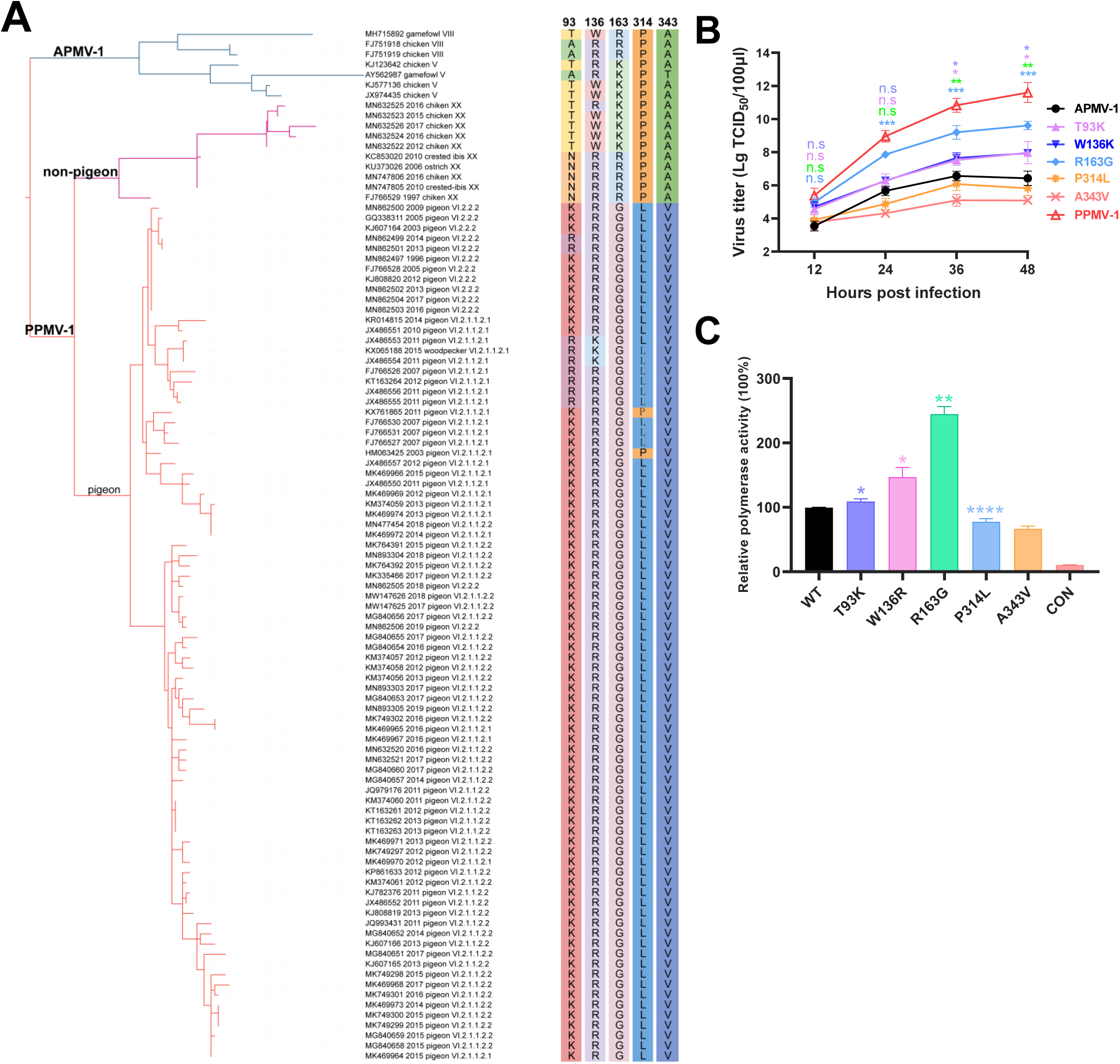
A single amino acid change in P protein switches host preference. **A.** Amino acid substitutions of P protein of the APMV-1 and PPMV-1 strains. The genotypes and hosts of PPMV-1 were labeled on the branches. **B.** Virus titers of PPMV-1 with single - amino acid mutations. Error bars represent the SD of the mean from one representative experiment three biological replicate samples and each experiment was repeated three times. *: P ≤ 0.05; **: P < 0.01; ***: P < 0.001; ****: P<0.0001; n.s: not significant (P > 0.05) by t test. * labeled by different colors correspond to different mutant sites. **C.** Polymerase activities of wild-type and mutants. The polymerase activity of each mutant strain was compared with wild type strain. Error bars represent the SD of the mean from one representative experiment three biological replicate samples and each experiment was repeated three times. *: P ≤ 0.05; **: P < 0.01; ***: P < 0.001; ****: P<0.0001; n.s: not significant (P > 0.05) by t test. * labeled by different colors correspond to different mutant sites.

**Figure 4.**
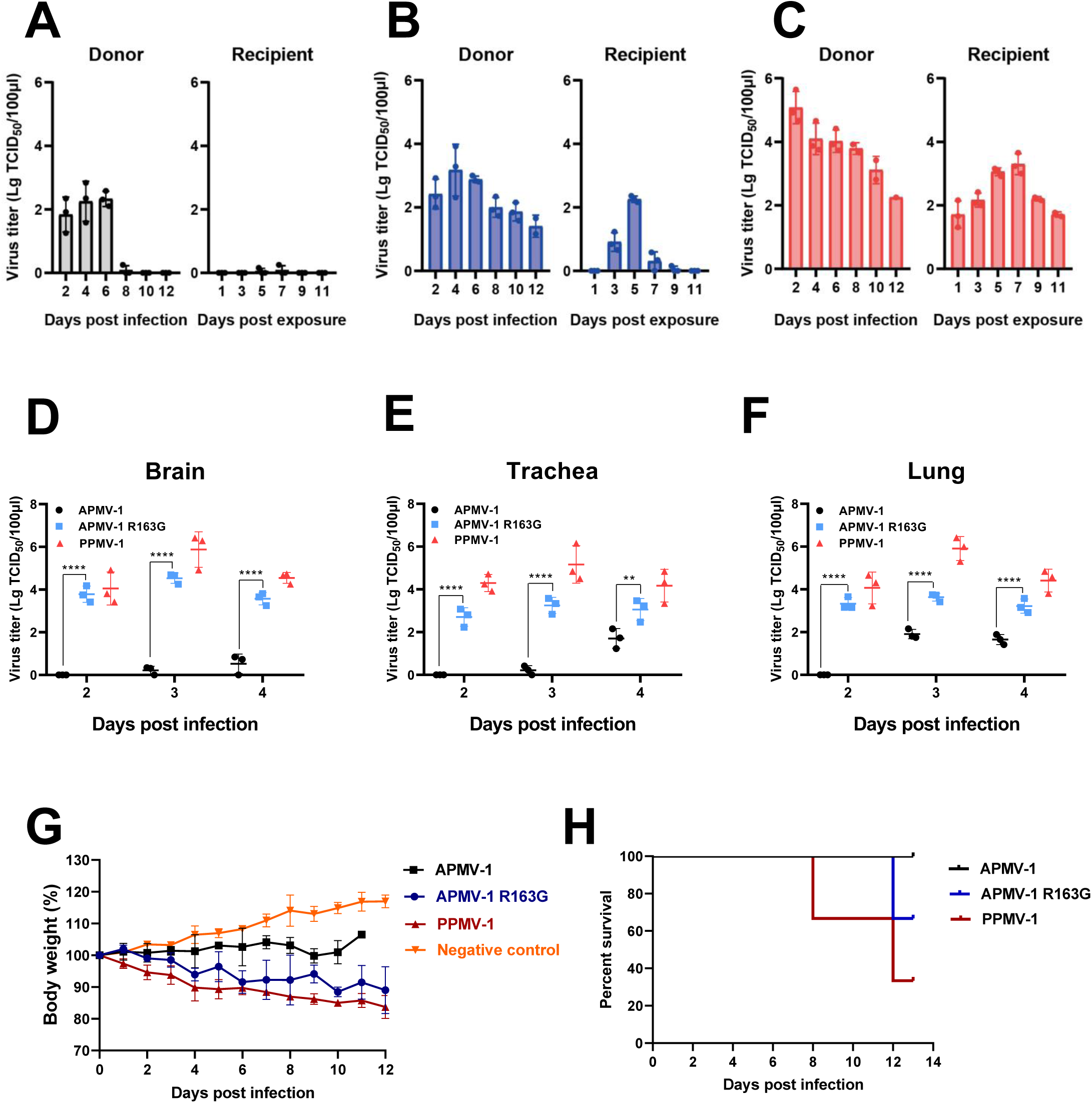
APMV-1 R163G increased pathogenicity and transmissibility in pigeons through increased degradation of host proteins. **A-C.** Contact transmissibility of APMV-1 **A.**, APMV-1 R163G **B.**, and PPMV-1 **C.** in pigeons. Each virus was tested in duplicate with a total of three donors and three direct contacts. **D-F.** Replication kinetics in pigeon brain **D.**, trachea **E.** and lung **F.**. Each data point represents one explant sample, and mean and SD are shown. Statistical differences were calculated by two-way ANOVA. *: P ≤ 0.05; **: P < 0.01; ***: P < 0.001; ****: P<0.0001; n.s: not significant (P > 0.05)

To validate whether the sites linked to adaptation, we incorporated P mutations separately in 93, 136, 163, 314, and 343 into the APMV-1 genome and analyzed the mutant virus. Pigeon fibroblast (PEF) cells were then infected with the mutant and wild type strains, and the virus titers were measured every 12 hours until 48 hours post infection. The growth curves of these APMV-1 mutants were variable. Mutant strains showed residues T93K, W136R, R163G significantly higher virus titers than wild type, especially site R163G, while the viral titer was reduced on the P314L and A343V mutants (fig. 3B). Further mini-genome activity tests also showed similar results (fig. 3C), which suggested the importance of these mutations in regulating the biological activity of polymerase. These results support the stepwise adaptation of PPMV-1 since its introduction from the APMV-1 through the three mutations on P protein.

We next focused on the three P protein amino acid substitutions of the virus in China. A phylogenetic analysis based on the complete P protein was conducted and found that Site 93 settled as K after multiple changes (T, N, R), Site 136 changed from W to R/K, while site 163 changed from R/K to G (supplementary fig. 5). Convergent evolution can be used to differentiate adaptation from neutral genetic variation on the basis of sequence data (Pepin et al, 2010). To verify whether the virus has undergone adaptive evolution, a convergent evolution analysis was performed in the United States. We found the three mutations are similar to those in China (I93K, R163G, beisides, 136R is consistent with the mutated site of the Chinese strain) (supplementary fig. 6).

### APMV-1 R163G Increases Transmission Efficiency and Pathogenicity in Pigeons

Since R163G is the most sufficient site to improve the replication capacity in PEF cells, we thus compared the contact transmission potential of APMV-1, APMV-1 R163G, and PPMV-1 viruses in 4 weeks old pigeons (the PPMV-1 isolate was used as positive control). The viral load of APMV-1 R163G group in the ranges of 1.15-3.91 lg TCID_50_, higher than those inoculated with APMV-1, with the viral load in the ranges of 1.25-2.83 (figs. 6B and 6C). In contact pigeons, APMV-1 R163G viral load in the ranges of 0.34-1.23 lg TCID_50_, higher than those exposed to APMV-1, with viral load in the ranges of 0.15-0.25 lg TCID_50_ (figs 6B and 6C). These results suggest that APMV-1 R163G virus showed better replication in inoculated donors and transmitted more rapidly to contact pigeons than the APMV-1 wild type isolates.

We found that APMV-1 R163G replicated to higher titers than APMV-1 at 1 day post-infection (dpi) in pigeon lung, trachea, and brain explants from 2 dpi to 4 dpi (figs. 6D, 6E and 6F). Moreover, pigeons infected with APMV-1 R163G displayed rapid weight loss than that infected with APMV-1 wild type (fig. 6G). One pigeon died on 12 dpi. As expected, the pigeons infected with APMV-1 all survived (fig. 6H).

**Figure 5.**
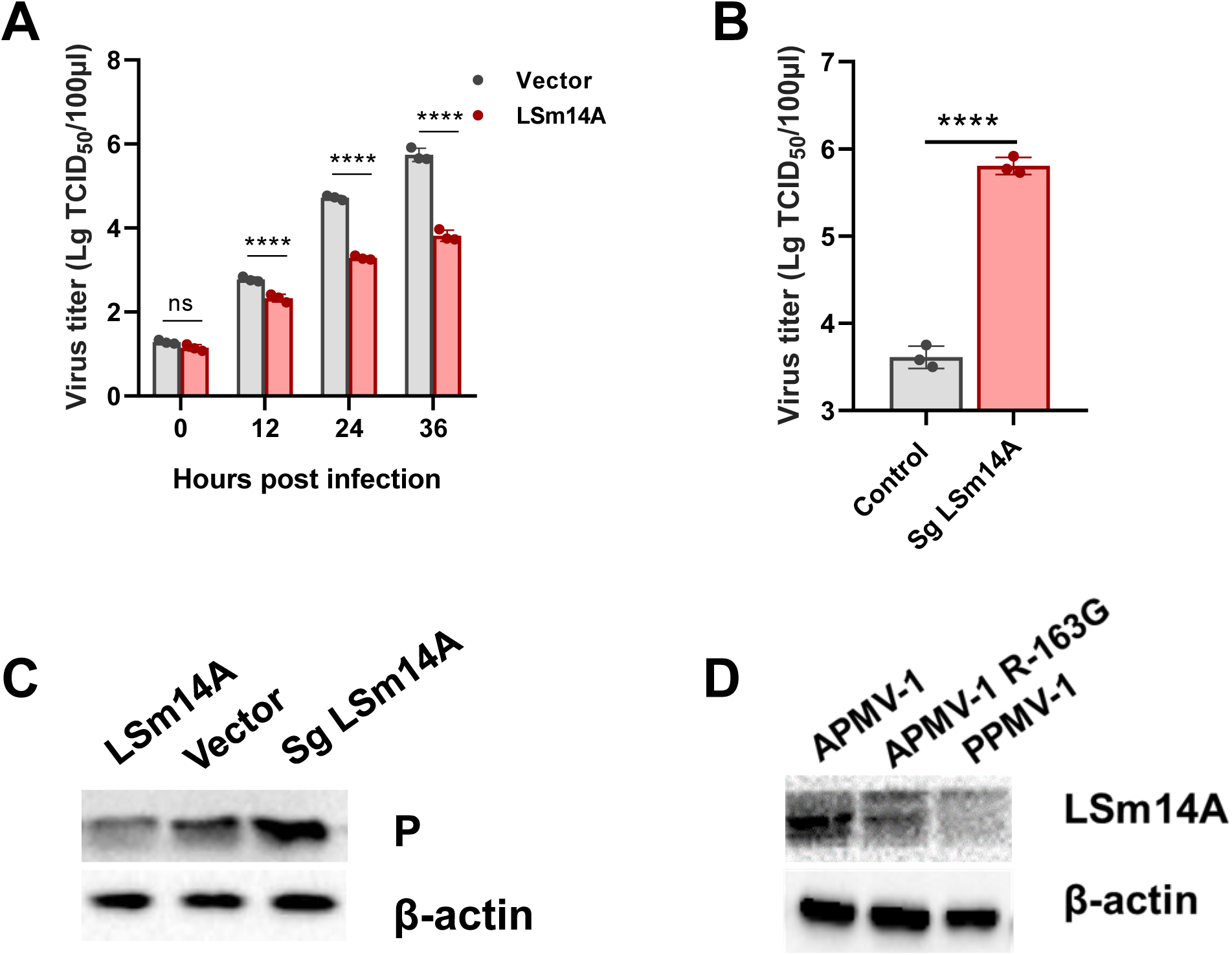
The mutation on P R163G can enhance the virus replication by increase degrade LSM14A protein. **A.** Inhibition of APMV-1 titer by overexpression of LSM14A. **B.** Enhancement of APMV-1 titer by CRISPER-Cas9 knockout of LSM14A. **C.** PRRSV replication is inhibited by LSM14A on protein level. **D.** The mutation on P R163G increases the ability to degrade LSM14A protein.

**Figure 6.**
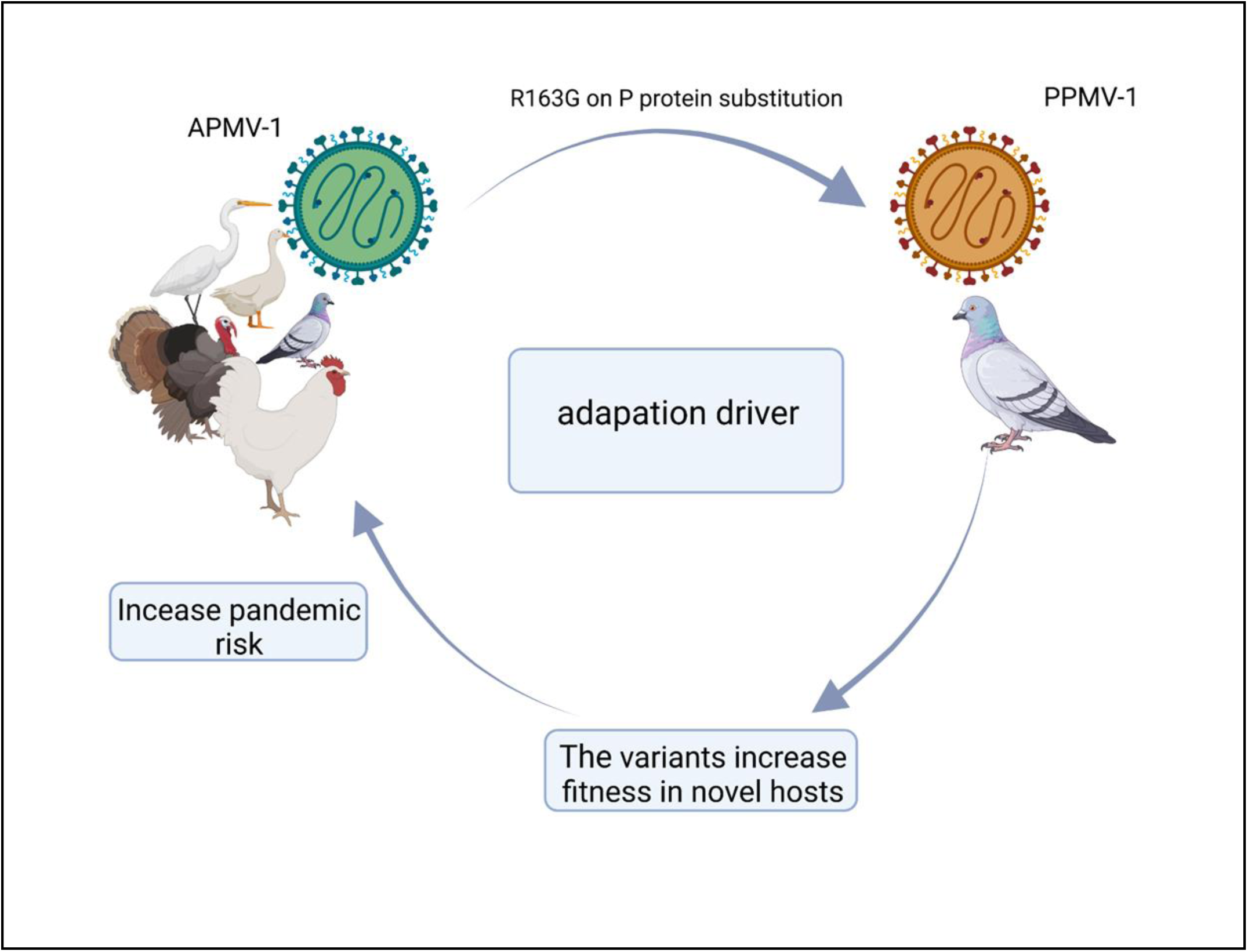
Schematic model to show the evolution from APMV-1 to PPMV-1. Host adaptation drives evolution of APMV-1 to PPMV-1. Three amino acid residues on P protein play an important role in host shift. The well-fit variants increasing the risk of a pandemic.

### APMV-1 R163G Enhances Its Replication Ability by Increasing Degradation of LSm14A

Since the P protein is associated with host shift, we set out to identify host cellular factors that associated with the P protein. We focused on LSm14A since it has the highest sum PEP score as identified by LC/MC. LSm14A is a key innate immunity component of processing body (P-body) that mediates interferon-β (IFN-β) signaling by viral RNA (Li et al, 2012). To confirm whether chicken LSM14A could inhibit APMV-1 replication, we overexpressed chicken LSm14A in DF-1 cells. The results showed that expression of LSm14A significantly inhibited APMV-1 replication (figure 7B). To further confirm that LSm14A could inhibit APMV-1 replication, a comparative analysis was carried out by CRISPER-Cas9 at later time points (36h) to examine whether the knockdown of LSm14A could enhance APMV-1 replication. The results showed that knockout of LSm14A significantly enhanced APMV-1 replication at 36 h post infection (fig. 7C). This result was also verified by Western blot (fig. 7D). The R163G mutation on P protein increased degradation of LSm14A (fig. 7E), thus it may facilitate escape from host immune responses.

## Materials method

### Ethics Statements

These animal studies were performed in strict accordance with the Guidelines for the Care and Use of Animals in Research, which are issued by the Institute of Zoology, Chinese Academy of Sciences (Approval Number IOZ12017).

### Cells, Viruses and animals

The pigeons were bought from a hatchery in Miyun District, Beijing, and certified by hemagglutination inhibition (HI) experiment to have no antibodies of NDV and AIV. These pigeons were housed in separate cage in an animal room under biosafety conditions with a suitable temperature and an adequate supply of food and water.

The APMV-1 strain F48E9 (MG456905) was kept in our laboratory, and the mutant viruses were produced from an infectious cDNA clone.

The PEF cells were isolated from 10-day-old pigeon embryos, and maintained in DMEM (Gibco, 11965–092) supplemented with 10% FBS and 1× penicillin, streptomycin.

### Bayesian Phylogenetic Analysis

The complete F gene (1662bp) sequences from PPMV-1 isolates were downloaded from Genebank and aligned using MEGA version 7. We selected the best fitting model using jModelTest v2.1.7 on the basis of the Akaike Information Criterion (AIC). Time-scaled phylogenetic analyses were conducted using an uncorrelated relaxed clock with the GMRF Bayesian tree and general time-reversible (GTR) model with gamma 4 substitution and invariant site model parameters jModelTest output. Molecular clock was calibrated under an uncorrelated relaxed clock grouped with different trees using BEAUti (v1.8.4). The Bayesian Markov Chain Monte Carlo (MCMC) chain length was 100,000,000 generations with sampling every 10,000 generations. Mixing was assessed using effective sample size (ESS) using Tracer (v1.6). The MCC tree at each iteration was generated by TreeAnnotator v.1.8.4, discarding first 10% of the chains as burn-in. The resulting MCC tree was visualized with FigTree software (v1.4.3) and edited with Adobe Instructor. Bayes factors (BFs) were used for posterior probability calculation.

### Bayesian Skyline Plot Construction

A coalescent-based Bayesian skyline plot was implemented using a piecewise-constant skyline model from the BEAST version 1.8.4 software. This plot was used to quantify contributions of potential predictors of PPMV-1 dispersal in China and the world.

### Discrete Phylogeography and Transmission between Hosts

To infer Bayes factors that are associated with PPMV-1 dispersal among countries and regions within China, we used Bayesian stochastic search variable selection (BSSVS) to determine the most probable locations of ancestral nodes in the phylogeny and the history and rates of lineage movement among locations (Lemey et al, 2009). Complete F gene sequences from China and the rest of the world were selected to analyze using the BSSVS method. Six candidate clocks with models were compared using a Path Sampling/Stepping-stone analysis in order to estimate the best-fitting demographic model for our dataset. We then used BF comparisons from the resulting marginal-likelihood estimates to select among the models. We found that the uncorrelated clock with constant model provided the best fit to our data (Supplementary Table1).

We inferred the global and within-China geographic origins of PPMV-1 and its significant dispersal routes between affected countries or regions using discrete-state ancestral reconstruction methods implemented in BEAST SPREAD3 version 0.9.6. We used BF > 3 as the significance threshold.

The results of BSSVS were summarized using spreaD3 v0.9.7.1 (Bielejec et al, 2011), a json file was generated to identify the routes of geographic diffusion and their associated Bayes factors and the spatio-temporal pathways of PPMV-1 were visualized by Echart (https://echarts.apache.org).

To model transmission between host species, chicken, pigeon, and wild birds were used as discrete traits. We combined the best fitting coalescent tree model and branch-rate prior as described above. The BSSVS method was used to identify significant migration routes and their directionality between hosts. Using the Markov-jump (MJ) method, the intensity of backward and forward transitions within discrete trait matrices was inferred as a proxy for the mean number of viral jumps between hosts.

### Selection Pressure Analysis of PPMV-1 Proteins

Sequences of the six PPMV-1 proteins were aligned using MEGA 7.0, respectively. After deleting terminators, we used datamonkey (http://www.datamonkey.org/) to analyze selection pressure. The BUSTED (Branch-site Unrestricted Statistical Test for Episodic Diversification) method was used. Selection intensity was expressed as dN/dS.

### Synonymous and Non-synonymous Substitution Rate Analysis

The synonymous and non-synonymous substitution rates were estimated using SNAP (Synonymous Non-synonymous Analysis Program) v2.1.1 (https://www.hiv.lanl.gov/content/sequence/SNAP/SNAP.html). The results are presented by cumulative dN/dS, with dN>dS indicative of positive selection.

### Statistical Analyses of the F and P proteins

The F and P protein amino acid sequences were type-entered into a database and analyzed using SPSS software version 20. Categorical variables were compared using the Pearson χ2 test, Fisher’s Exact Test or Linear-by-Linear. Comparison group pair-wise P-values were corrected using the Bonferroni method. P < 0.05 were considered statistically significant.

### Mutations Linking Host Shift via treeWAS

We selected 101 viral sequences that infect chickens or pigeons. The protein sequences of the F or P gene were aligned with MUSCLE v3.8.31 (Edgar 2004), and the codon alignments were made based on the protein alignment with RevTrans (Wernersson and Pedersen 2003). RAxMLv8.2.12 (Stamatakis 2014) was used to build the maximum likelihood phylogenetic tree of aligned codon sequences with the parameters ‘-p 1234 -m GTRCAT’. Based on codon alignments and phylogenetic trees, we screened possible host-related loci using treeWAS (Collins and Didelot 2018).

### Virus Rescue and Site Mutation

A reverse genetics system for the generation of recombinant viruses of the APMV-1 strain as previous research (Dortmans et al, 2009).

The mutations on P protein were conducted with Mut Express II Fast Mutagenesis Kit V2 (Vazyme, C214), and the mutant and vector linearization primers were designed on Vazyme website (https://crm.vazyme.com/cetool/singlepoint.html). The primer sequences list on Supplementary Table2.

### Viral Replication Kinetics in Vitro

PEF cells in 96-well plates were inoculated mutant and wild strains at an MOI of 0.01. Supernatants were collected at 12, 24, 36, 48 hpi and virus titers were determined via limiting dilution in PEF cells using the approach of Reed and Muench and expressed as the 50 % tissue culture infective dose (TCID_50_).

### Mini-genome PPMV-1 System and Dual Luciferase Assay

To detect whether mutations in the P protein influenced polymerase activity, a mini-genome plasmid was constructed. 3’- and 5’-UTRs derived from KJ607169, the firefly luciferase gene, was reverse complementary cloned into pOK12, followed by the hepatitis delta virus ribozyme and the terminator. BSR T7 cells were seeded in 6-well culture plates and co-transfected with plasmids each expressing NP (2.5 μg), P mutated or WT (1.25 μg), and L (1.25 μg) proteins, as well as a firefly luciferase reporter gene (RLuc, 20 ng) with a Renilla luciferase expression plasmid (phRL-TK) as an internal control. The washed cells in PBS twice at 24 h after transfection, added 200 μl cell lysis buffer to each well, and shocked for 10 min. 20 μl of lysate from each well was used for dual the luciferase assay using a commercial kit (Vazyme, DL101-01). The relative luciferase activities were defined as the ratio of FLuc to RLuc according to the manufacturer’s instructions (Dortmans et al,, 2010). Three separate experiments were performed, with luciferase expression measured in triplicate in each experiment.

### Compare the Pathogenicity of APMV-1, APMV-1-R163G and PPMV-1

A total of 30 male and female pigeons at 4 weeks old, with approximately equal body weight of −10-10% were used in our study. These pigeons were randomly divided into six groups, and housed in 12 h dark and 12 h light environments with a suitable temperature and an adequate supply of food and water. Three groups of four-week-old pigeons of six birds each inoculated with 10^6^ EID_50_ of virus in a volume of 200 μl via intranasal route. Another three groups of pigeons representing contact groups were inoculated with 200 μl PBS via the same routes and co-housed with the three inoculated groups of pigeons to monitor contact transmission between pigeons after 24 hours after inoculation. The negative control group was inoculated with 200 μl via intranasal route. Three pigeons of each group were euthanized at 3 dpi, and the remaining birds were monitored for illness and mortality for 14 days.

### Tissue Distribution

At 3 days post-inoculation (dpi), three from the four groups were euthanized to analyze viral replication in the brain, lung, and trachea. The virus was isolated from tissues of the same weight, and TCID_50_ in DF-1 cells (chicken fibroblast cell line) was used to estimate viral loads of the four groups, 3 × 10^4^ DF-1 cells were seeded in 96-well plate with five repetitions 1 day before infection. Twenty-four hours later, the cells were infected with different dilutions of the virus for 1 h at 37°C with shaking every 12 h and confirmed by the hemagglutination assay. TCID_50_ was calculated using the Reed-Muench method. Data were analyzed using Prism (v.5.01) software. Statistical significance was set at a P-value of <0.05 (Two-way ANOVA).

### Compare the Transmissibility of APMV-1, APMV-1-R163G and PPMV-1

At 1, 3, 5, 7, 9, 11 and 13 dpi, oropharyngeal swabs were collected from all pigeons and placed in tubes with phosphate-buffered saline solution and 2% fetal bovine serum and stored at −80°C until RNA extraction.

### Identification of Host Proteins Related to the P protein

P-His was ligated into the Vector pFastDual, and verified by Western blot analysis (data not shown). Lysates were prepared from Sf9 cells that were infected with baculovirus expressing P-His protein; uninfected Sf9 cells were used as a control. Affinity purification by Nickel column purification (Smart lifesciences) was directed against the His tag; therefore only host proteins that associated with P were precipitated by this method (Supplementary fig. S6A). The samples were separated using a 10% SDS-PAGE gel, followed by Protein bands were visualized using Coomassie brilliant blue staining (Supplementary fig. S6B). The indicated protein-containing band was cut out for mass spectrometry (Supplementary Table3). LSm14A scores highest. We then used co-immunoprecipitation confirmed the interaction of LSm14A with P protein in DF-1 cells.

### Knockout and Overexpression

Chicken LSM14A (Gene ID: 415781) was cloned from cDNA of the DF-1 cells, using the following primer sequences: LSM14A-F: taagcttggtaccgagctcggatcATGAGCGGGGGGACGCCCTACATC, LSM14A-R: cactgtgctggatatctgcagaattcCTATGCAGCAACTTTGTTGTCTTTCCTATATTCAAAGTCAGCAAACTCCC. The purified PCR product was digested with BamH I and EcoR I, and inserted into the eukaryotic expression vector pcDNA 3.1 (vector).

We designed two sgRNA for CRISPR knockout of LSM14A using design.synthego.com: LSMsg-F1, CACCGCTGGCCAAGGTTCGTTCCTT; LSMsg-R1, AAACAAGGAACGAACCTTGGCCAGC; LSMsg-F2, CACCGATACCACTGCGTCCTAATCG; LSMsg-R2, AAACCGATTAGGACGCAGTGGTATC. The sgRNA were then cloned into Lenti CRISPER plasmid for subsequent transfection into 293T cells.

### Western Blot Analysis

Protein was extracted from cells in Radioimmunoprecipitation assay (RIPA) lysis buffer containing 1× complete Protease Inhibitor Cocktail (bimake). Samples were analyzed by SDS-PAGE and followed by electrophoretic transfer to polyvinylidene fluoride membranes, which were then blocked and incubated with primary antibodies. The following antibodies were used in the experiments: anti-LSm14A (dilution 1:1000), anti-P (dilution 1:2000), β-actin (dilution 1:5000).

## Disscussion

PPMV-1, an APMV-1 variant, establishes a species-specific relationship with its new host during the evolution process. In this study, we provide unique insights into the drive factor of PPMV-1 jumping from multiple birds to pigeons via a series of analyses. We clarified adaptation driver is the main factor on host shift rather than ecological driver. P protein plays an important role in host shift by select pressure and treeWAS analysis. We demonstrate through molecular biology experiments the variants increase fitness in novel hosts via increasing degradation sensor-LSm14A. The well-fit variants replicate efficiently in the new host, increasing the chance of spillover to other hosts, thereby increasing the risk of a pandemic (fig. 5).

The evolutionary analysis on a global scale shows the virus in some countries compartmentalizing the genotypes within geographically discrete, possibly due to relatively low country-to-country transmission, especially in China and the United States. Isolates from countries with frequent communication typically cluster together. Europe was the epicenter of global PPMV-1 dissemination, it might because of the movement of pigeons through commercial trade (Chong et al, 2013; Hicks et al, 2019). In China, the initially dominant subtype is gradually replaced by a new subtype and repeat the process, and formed a ladder-like PPMV-1 phylogeny. This evolution mode implies the presence of selection pressure, probably driven by host immune escape (Volz and Frost 2013). Regions within the country share subtypes without observable differentiation. Viral spread occurred most frequently from south to north China, possibly because these regions are on the border between the Central Asian and East Asian/Australian migratory flyway (Olsen et al, 2006). Industry practices that involve transporting birds born in the South to be raised and bred in the North, also likely contribute to this pattern of PPMV-1 spread. Overall, these results suggest that animal exchange is an important contributor to the dissemination of PPMV-1 subtypes.

The proportion of pigeons infected by PPMV-1 increased with its evolution, and pigeons are the most susceptible host species to this virus (Smietanka et al, 2014; Mayahi et al, 2017; Zhan et al, 2020), indicating that the PPMV-1 evolution trajectory is host-specific rather than host expansion. Since its emergence, PPMV-1 has continued to adapt for pigeon infection, resulting in the emergence of variants with unique genetic profiles. Viral adaptation to new hosts primarily manifest as amino acid substitutions that allows more efficient virus cell entry in the novel host (Ito et al, 1998), blocks interactions with detrimental host proteins (Mangeat et al, 2003; Stremlau et al, 2004) or promotes escape from both the new and the old host’s immune responses (Wei et al, 2003; DJ et al, 2004). The probability of a host shift will also depend on the number of mutations required to adapt to novel hosts. Host shifts are often associated with changes in the viral polymerase complex (Gabriel et al, 2005; Ackermann et al, 2007). Our treeWAS analysis shows five non-synonymous on P protein (a member of polymerase complex) may associate with host shift. In some cases, adaptation to a novel host relies on specific mutations that arise repeatedly whenever a pathogen switches to a given host (Longdon et al, 2018). In this study, the three selected PPMV-1 P protein sites (T93K, W136R, R163G) underlie the APMV-1 evolution to PPMV-1. A previous research indicated that Genotype VIII evolved as Genotype XIX and V, then evolved Genotype XX PPMV-1 (Dimitrov et al, 2019), thus Genotype VIII and V were used to represent APMV-1 for analysis. From the evolutionary trends, we can clear see the mutations occur when host shift rather than genotype shift. Each amino acid substitutions, whether affects adaptation are measured by a mutational fitness effect. Five mutations on the reconstituted P protein show that single mutations at three sites could enhance the replication of APMV-1 in pigeon embryo fibroblasts, and the mutation R163G is the most effective. Importantly, R163G facilitated the transmission and pathogenicity in pigeons. LSm14A is known to be involved in antiviral signaling pathway, and its transcriptional levels of the spleen tissues remained high after F48E9 (high virulent) and P3 (moderate virulent) infection (Tian et al, 2019). Degradation of LSm14A may help viruses evade host defenses (Saeed et al, 2020), thus APMV-1 R163G is more pathogenic and transmissible than APMV-1.

The single mutation on virus could switch the host preference, which is in line with previously reported findings. For example, Venezuelan equine encephalitis virus to replicate efficiently in horses when it switched from rodents in the early 1990s (Anishchenko et al, 2006), the single mutation in the AIV HA protein that changed receptor binding preferences from α-2, 6 to α-2, 3 (Nicholls et al, 2007), and the epidemic in humans associated with a CHIKV strain carrying a single mutation in the envelope protein gene (Tsetsarkin et al, 2007). Amino acids 136 and 163 on the P protein are located in a hypervariable motif at residues 135-165 near the RNA editing site, adjacent to the cysteine enriched region of the V protein that constitutes the zinc finger structure. We speculate that the K163G mutation changed PPMV-1 host preference via alterations in alkaline acid similar to PB2 627 of AIV (Lee et al, 2019; Liu et al, 2019). The mutation on P protein causes a mutation (E163R) on the V protein. The C-terminal region of the APMV-1 V protein exerts IFN antagonistic activity (Park et al, 2003), of which the residue 163 located. Likewise, the mutation changed the amino acid charge of the V protein.

Our findings inferred that PPMV-1 may have originated from a variety of birds, the intermediate amplifying and transmitting host, which consistent with previous findings (Chong et al, 2013). Our BSSVS analysis indicated that the virus tends to spill over from the susceptible host, but seldom comes back, which consistent with previous findings (Xiang et al, 2020; Chen et al, 2021). We inferred that the virus accumulates in the susceptible host due to its efficient replication capabilities, with genotypes exhibiting high titers more likely to spill over into other hosts.

PPMV-1 are not easy to infect species other than their natural host, whereas the occasional cross-species transmission to other flocks or animals has been documented, several most spill-overs result in a very severe disease in the new host, causes a pandemic. Some studies suggested that multiple ND outbreaks, especially with the third pandemic in Great Britain in 1984, were initiated by PPMV-1 that spread from pigeons to chickens (Alexander 1988; Aldous et al, 2004). Subsequently, PPMV-1 was responsible for other chicken ND outbreaks worldwide (Werner et al, 1999; Kommers et al, 2001; Capua et al, 2002; Aldous et al, 2004; Liu et al, 2006). The PPMV-1 can cause severe pneumonia and death in humans, however, we can’t obtain the P protein sequences from these reaches. We can infer the Site 163 is G from the subtype, thus human infection may be an accidental event. We can use the degradation ability of LSm14A for preliminary explored infectivity of a PPMV-1 strain.

USA, another country has special PPMV-1 subtypes, VI.2.1 and VI.2.1.1.1, were displayed in the evolutionary tree. Due to the limited number of PPMV-1 strains from USA, we could not make robust inferences of evolutionary forces acting on the P protein. These subtypes are distinct from those found in China but contain the similar mutations. These parallel mutations provide compelling evidence that these genetic changes are adaptive, with the same mutations evolving independently in response to natural selection. Only a strong selective force is likely to cause multiple occurrences of the same genetic make up originating from different starting points. Taken together with the data from the APMV-1 virus, these results point to repeated evolution of the same P protein sequences underlying host shifts, strongly suggesting that the establishment of a new genotype will depend on ecological and host species traits, and adaptive evolution is the main evolution driving force.

In present study, we document a stepwise adaptation of PPMV-1 to its pigeon hosts involving multiple amino acid changes that appear essential for the successful host shift and prolonged transmission. Our research highlights the importance of the amino acid substitutions on P protein as a marker of both PPMV-1 origin and species-specific polymerase function. Strictly host-specific viruses can cause epidemics once cross-species transmission occurs. The emergence of PPMV-1 demonstrates how a virus can successfully cross species barriers and become established as an epidemically spreading pathogen in a new host.

## Acknowledgments

The work was supported by Strategic Priority Research Program of the Chinese Academy of Sciences (XDA19050204), National Forestry and Grassland Administration of China, Beijing Wildlife Rescue Center of China, and National Key R&D Program of China.

## Author Contributions

H.C., H.H. and J.L designed the project. H.C., S.F. and X.T performed the experiments. H.H. obtained the funding for the project. H.C., W.S., S.F., X.T, C.X. Z.W and B.W. analyzed the sequencing data. H.C., S.J., Y.W. and Y. X coordinated the animal experiment. H.C., S.F, Z.W, B. H and G.L wrote the manuscript with the help of all authors.

